# NEFFy: A Versatile Tool for Computing the Number of Effective Sequences

**DOI:** 10.1101/2024.12.01.625733

**Authors:** Maryam Haghani, Debswapna Bhattacharya, T. M. Murali

## Abstract

**Summary:** A Multiple Sequence Alignment (MSA) contains fundamental evolutionary information that is useful in the prediction of structure and function of proteins and nucleic acids. The “Number of Effective Sequences” (NEFF) quantifies the diversity of sequences of an MSA. Several tools can compute the NEFF of an MSA, each offering various options. NEFFy is the first software package to integrate all these options and calculate NEFF across diverse MSA formats for proteins, RNAs, and DNAs. It surpasses existing tools in functionality without compromising computational efficiency and scalability. NEFFy also offers per-residue NEFF calculation and supports NEFF computation for MSAs of multimeric proteins, with the capability to be extended to nucleic acids (DNA and RNA).

**Availability and Implementation:** NEFFy is released as open-source software under the GNU General Public License v3.0. The source code in C++ and a Python wrapper are available on GitHub at https://github.com/Maryam-Haghani/NEFFy. To ensure users can fully leverage these capabilities, comprehensive documentation and examples are provided at https://Maryam-Haghani.github.io/NEFFy

## Introduction

A Multiple Sequence Alignment (MSA) organizes a set of similar sequences by introducing gaps to ensure that all sequences are of the same length, *l*. The MSA contains each sequence in a row with *l* columns or positions, where each residue or gap occupies a distinct position. Computing an MSA involves maximizing the similarity in each position across the rows while minimizing the number of gaps. By uncovering evolutionarily conserved sequence patterns, regions of similarity that cannot be identified from a single sequence alone, MSAs are used in applications including contact map prediction [1, 2, 3, 4], RNA and protein structure prediction [5, 6, 7, 8, 9], and protein function annotations [10, 11, 12].

In recent applications, construction of an MSA starts with a query sequence of interest. The process involves searching databases to find sequences similar to the query and aligning them to it. Recent advancements in DNA/RNA sequencing technology have expanded public databases, enabling the generation of MSAs with high sequence diversity [13, 14]. Such MSAs are generally believed to provide richer evolutionary and coevolutionary insights, and they can thereby enhance the effectiveness of models utilizing them for downstream tasks [9]. However, since MSAs can contain redundant sequences, the number of sequences by itself may not be an accurate reflection of their diversity. The concept of “Number of Effective Sequences”, NEFF, addresses this redundancy and assesses the quality of an MSA. Higher NEFF values often indicate a more diverse and informative MSA, leading to enhanced accuracy in predicting contact maps and the tertiary structures of proteins or RNA molecules [15, 16]. For example, the accuracy of AlphaFold declines substantially when the NEFF value is below approximately 30 [5]. Additionally, for RNA structure prediction models such as trRosettaRNA, which utilize MSAs of RNAs as their input, prediction accuracy is correlated with NEFF [7], and for high-quality MSAs, these models can outperform other methods [17].

We introduce NEFFy, a fast and dedicated standalone tool for NEFF calculation. NEFFy is uniquely equipped to parse MSAs and calculate NEFF across a wide range of MSA formats for protein and nucleic acid sequences. It integrates all the features from earlier NEFF tools (see Table 1) and offers a set of new functionalities. NEFFy is developed in C++ for optimal performance and is also provided as a Python library that wraps the C++ executables. This approach enables seamless integration into Python-based workflows, simplifying use for a wider audience while preserving efficiency.

**Table 1:** An overview of NEFFy and other tools that incorporate NEFF calculation, highlighting their respective features (refer to Supplementary Section S2.1 for detailed information on each feature). **Language**: The programming language used to implement each tool. **MSA Format**: Formats used to represent aligned sequences in an MSA. **Similarity:** Methods for determining sequence similarity between pairs of sequences within the MSA. **Query Gaps**: Handling query gaps—defined as “gaps aligned to insertions”—can be customized during NEFF calculation based on user preference: either by removing them along with the corresponding positions in the aligned sequences or by keeping them intact. ^*^Conkit can manage these gaps for the a3m format by offering two distinct options: a3m-inserts and a3m. **Gappy position:** Inspired by Gremlin, this option addresses positions with a gap frequency exceeding the desired gap threshold. **Alphabet**: Alphabet used to represent a biological sequence. Of the tools mentioned, RaptorX and Conkit do not explicitly define an alphabet, while DeepMSA and rMSA use the protein alphabet. In contrast, Gremlin supports both protein and RNA sequences (implicitly including DNA). NEFFy accommodates proteins, RNAs, and DNAs. **Non-standard Residues**: Residues outside the standard set of biological sequence residues are handled differently across tools. RaptorX and Conkit do not support these residues, while rMSA and Gremlin treat them as gaps. DeepMSA considers them as standard symbols in its symmetric version but treats them as gaps when calculating similarity cutoff in its asymmetric version. NEFFy offers users the flexibility to customize the handling of these residues according to their preference. **Validation**: Indicates if validation is provided for the MSA file before the NEFF calculation.

| Tool | Language | MSA Format |  |  |  |  |  |  | Similarity |  | Query Gaps | Gappy Position | Multi Alphabet | Non-Standard Residues | Validtion |
| --- | --- | --- | --- | --- | --- | --- | --- | --- | --- | --- | --- | --- | --- | --- | --- |
|  |  | a2m | a3m | sto | aln | fasta | clustal | pfam | sym. | asym. |  |  |  |  |  |
| RaptorX [18] | Python | ✓ | ✓ | ✗ | ✗ | ✗ | ✗ | ✗ | ✗ | ✓ | ✗ | ✗ | ✗ | ✗ | ✗ |
| Conkit [19] | Cython | ✓ | ✓ | ✓ | ✗ | ✓ | ✓ | ✗ | ✓ | ✗ | ✗* | ✗ | ✗ | ✗ | ✗ |
| DeepMSA [14] | C++ | ✗ | ✗ | ✗ | ✓ | ✗ | ✗ | ✗ | ✓ | ✓ | ✗ | ✗ | ✗ | ✗ | ✗ |
| Gremlin [20] | C++ | ✗ | ✗ | ✗ | ✓ | ✓ | ✗ | ✗ | ✓ | ✗ | ✗ | ✓ | ✓ | ✗ | ✗ |
| rMSA [21] | C++ | ✓ | ✗ | ✗ | ✓ | ✓ | ✗ | ✗ | ✓ | ✗ | ✗ | ✗ | ✗ | ✗ | ✗ |
| NEFFy | C++ & Python | ✓ | ✓ | ✓ | ✓ | ✓ | ✓ | ✓ | ✓ | ✓ | ✓ | ✓ | ✓ | ✓ | ✓ |

## Method

The NEFF of an MSA *M* can be formulated as

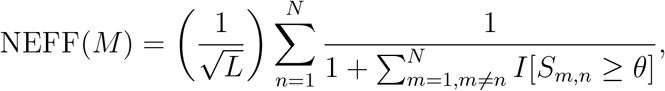

where *L* represents the length of the query sequence, *N* denotes the number of sequences in *M, S*_*m,n*_ indicates the amount of sequence similarity between *m*-th and *n*-th sequences, *θ* is the similarity threshold, and *I*[] is the Iverson bracket, meaning that *I*[*S*_*m,n*_ ≥ *θ*] equals 1 if *S*_*m,n*_ *≥ θ*, and 0 otherwise. Note that 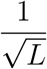 is a normalization factor, which is a commonly used approach, though other normalization options are also available. The NEFF value represents the normalized sum of the weights of the sequences in the MSA. If sequence *i* is similar to *n*_*i*_ other sequences (including itself), its weight is 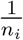. This approach for calculating NEFF is widely employed in many contact prediction and MSA generation methods [4, 14, 16, 22].

Aligning with the given formulation, several tools incorporate NEFF calculation as part of their functionality [14, 18, 19, 20, 21]. Additional details about each is provided in Supplementary Section S3 of the supplementary file. However, none of these tools are exclusively designed for NEFF calculation and they typically offer NEFF as required for their primary tasks. Consequently, they do not offer the full range of NEFF-related capabilities. Table 1 compares these tools with NEFFy in terms of the features they provide. It highlights that NEFFy is the only solution compatible with a wide range of MSA formats and supports all the features provided by the other tools. Moreover, NEFFy introduces several new features:

1. **NEFF calculation for multiple MSAs**: Accepts multiple MSA input files, combines their sequences in the specified order, removes duplicates, and calculates the NEFF for the resulting merged MSA.
2. **Per-residue (column-wise) NEFF calculation**: Calculates NEFF for each position in the MSA by summing the weights of sequences containing a residue (i.e., non-gap character) at that position.
3. **NEFF for multimeric MSAs**: Enables users to specify the multimeric format associated with the MSA, allowing NEFFy to calculate NEFF for all components.
4. **MSA format conversion**: Converts MSA formats while preserving sequence integrity and annotations, with no need for user intervention.
5. **MSA validation**: Ensures that the input follows the specified format and contains only residues from the permitted character set for the given alphabet.

Supplementary Section S2 contains detailed information about each feature.

## Results

We conducted an experiment using the CASP15 dataset, which consists of 93 targets, to compare the reliability and efficiency of NEFFy with other tools. To generate MSA files for these targets, we ran AlphaFold 2.3 locally, utilizing its default MSA generation pipeline. This process resulted in three MSA files per target, sourced from the Uniref90, Mgnify, and BFD datasets. In total, we generated 279 MSA files in `STO` and `A3M` formats. Given that certain tools do not support NEFF calculation in these formats, we employed NEFFy’s built-in converter to convert the files into formats compatible with each tool. We then calculated the NEFF value for each of these MSAs. To do this, we ran NEFFy using the options specified by each tool and also used each tool directly to calculate NEFF. The results showed that NEFFy consistently produced NEFF values identical to those of `DeepMSA` and highly similar to those from other tools (Supplementary Section S4.1).

To assess the computational efficiency of NEFFy relative to other tools, we used the MSA files from the previous analysis and recorded the execution time for each tool (Supplementary Section S4.2). NEFFy achieves the same efficiency as `DeepMSA` and `Gremlin` and significantly outperforms other tools. This makes it particularly ideal for processing deep MSAs of long query sequences (Supplementary Figure S6).

To evaluate the scalability of NEFFy with respect to MSA depth, we conducted an analysis by progressively increasing the MSA depth and measuring the corresponding execution times (Supplementary Section S4.3). The results show that NEFFy, along with `DeepMSA` and `Gremlin`, exhibits relatively constant execution times across all depths, demonstrating superior scalability and minimal sensitivity to increasing MSA depth. This indicates these methods are well-suited for efficiently handling deeper MSAs. In contrast, `RaptorX` and `Conkit` show a substantial increase in execution time as the depth increases, indicating that their performance is more sensitive to larger MSA depths and may struggle with higher computational loads at higher depths (Supplementary Figure S7). These findings provide valuable insights for selecting appropriate tools for MSA-based workflows depending on the computational resources available and the required scalability.

As a case study on multi-domain proteins, we explored the relationship between the NEFF values of individual domains and those of entire protein chains. Using 19 multi-domain proteins from the CASP15 dataset, we calculated NEFF values for the MSA of each domain and the entire target, using two separate sets of MSAs—one generated by AlphaFold [5] and the other by RoseTTAFold [6] (Supplementary Section S4.4). To compare the relative NEFF values, we generated Grishin plots [23], which display the correlation between the weighted sum of NEFF values for individual domains (y-axis) and the NEFF values for the full chain (x-axis). Our results showed that individual domains tend to have higher NEFF values than the complete protein chains in both the AlphaFold and RoseTTAFold-generated MSAs (Supplementary Figure S8).

In a recent study, Moussad et al. [24] examined the prediction accuracies of individual domains and their overall packing in tertiary structures generated by AlphaFold and RoseTTAFold models for these 19 multi-domain targets using the Global Distance Test (GDT-TS) metric designed to compare the similarity between predicted and reference structures [25]. They found that individual domains were predicted with high accuracy, while the overall packing of these domains was less accurate. Our NEFF analysis for the MSAs used as input for these multi-domain targets reflects similar patterns, suggesting that the higher NEFF values observed for MSAs of individual domains may contribute to the improved accuracy of their predicted structures.

## Conclusion

We present NEFFy, a fast and comprehensive tool for calculating the “Number of Effective Sequences” (NEFF) for a Multiple Sequence Alignment (MSA). By incorporating features from tools that already include this capability in their workflows and adding new functionality, NEFFy provides an efficient and versatile solution for NEFF calculation. Furthermore, NEFFy’s ability to convert and validate various MSA formats further enhances its usefulness. Additionally, NEFFy’s implementation as a Python library significantly broadens its accessibility and usability. Our experiments demonstrated that NEFFy is highly consistent with existing tools and is efficient, making it a valuable addition to the bioinformatics toolkit for processing MSAs of proteins and nucleic acids. As foundational models continue to evolve [26] and generate synthetic biological sequences, NEFFycan also be used to evaluate the effectiveness of evolutionary sequence search and alignment algorithms for proteins and nucleic acids (DNA and RNA) in comparison to synthetic sequences generated by these models.

## Supporting information

Supplementary Excel files containing NEFF values

## Data Availability

The MSA files used for NEFF analysis of reliability and scalability are hosted on Zenodo at https://zenodo.org/records/14199710, while those for multi-domain analysis can be found at https://zenodo.org/records/7682977.

