## Supplementary file for "NEFFy: A Versatile Tool for Computing the Number of Effective Sequences"

Table S1: Summary of Supplementary Excel Files and Sheets: This table provides an overview of each supplementary Excel file, detailing the sheet names, descriptions, and corresponding sections in the paper.

| **File Name** | **Sheet Name** | **Description** | **Notes** |
|---|---|---|---|
| Supplementary File S1.xlsx | CASP15 NEFF | NEFF values generated by running `NEFFy` and other tools on CASP15 dataset MSAs. | Related to S4.1 (first and second paragraphs). |
| Supplementary File S1.xlsx | `rMSA` NEFF | NEFF values generated by running `NEFFy` and `rMSA` on `rMSA` dataset MSAs. | Related to S4.1 (third paragraph). |
| Supplementary File S2.xlsx | Tool Timing (Average) | Average execution time for each tool based on 5 runs on CASP15 data. | Related to S4.2. |
| Supplementary File S2.xlsx | Scalability (Average) | Average execution time for each tool based on 5 runs across 36 CASP15 MSAs with varying depths. | Related to S4.3. |
| Supplementary File S3.xlsx | Multi-Domain Analysis | NEFF values for MSAs of entire proteins and individual domains in multi-domain CASP15 targets. | Related to S4.4. |

### S1 Introduction

NEFFy is a versatile and efficient tool for calculating the number of effective sequences (NEFF) across multiple MSA formats for various types of biological sequences.

A comprehensive user guide, including installation instructions and usage examples, can be found in the documentation at https://maryam-haghani.github.io/NEFFy/.

### S2 Options and Features

NEFFy is designed to seamlessly incorporate all NEFF calculation options provided by existing tools, as outlined in the subsections below.

#### S2.1 Integration of Existing Features

##### S2.1.1 Sequence Similarity

When determining the cutoff for sequence similarity, two approaches can be used: **1.** symmetric and **2.** asymmetric. In the symmetric approach, the similarity cutoff is uniform across all sequences in the MSA and is calculated as the product of sequence length and the given similarity threshold. Sequences are considered similar if the number of position-wise matches between them exceeds this cutoff. In contrast, the asymmetric approach only considers the non-gap positions in each sequence when calculating the number of position-wise matches, resulting in a variable similarity cutoff for each sequence. This leads to an asymmetric evaluation of sequence pairs. The default approach in NEFFy is symmetric, but users can switch to the asymmetric approach by setting is_symmetric to false.
While NEFFyRaptorX exclusively supports the asymmetric method and Conkit, Gremlin, and rMSA support only the symmetric method, only DeepMSA and NEFFy are capable of handling both approaches.

The pseudo code for computing sequence weights is outlined in algorithm 1.

**Algorithm 1** Sequence Weight Calculation

**procedure** COMPUTESEQUENCEWEIGHT($sequences$, $similarity\_threshold$)
  $sequence\_weight \leftarrow []$
  **if** $symmetric$ **then**
    $sequence\_cutoff \leftarrow length \times similarity\_threshold$
  **else**
    **for** each sequence $s$ in sequences **do**
      $non\_gap[s] \leftarrow$ non-gap positions of sequence $s$, considering value of $non\_standard\_option$
      $non\_gap\_count[s] \leftarrow$ number of non_gap positions
      $sequence\_cutoff[s] \leftarrow non\_gap\_count[s] \times similarity\_threshold$
    **end for**
  **end if**
  Initialize $similar\_sequences$ with all 1s (for the sequence, itself)
  **for** any sequence $s$ in sequences **do**
    **if** $symmetric$ **then**
      $similar\_sequences[s] \leftarrow$ count of sequences whose position-wise matches with sequence $s \geq sequence\_cutoff$
    **else**
      $similar\_sequences[s] \leftarrow$ count of sequences whose position-wise matches with non-gap positions of sequence $s \geq sequence\_cutoff[s]$
    **end if**
  **end for**
  **for** any sequence $s$ in sequences **do**
    $sequence\_weight[s] \leftarrow \frac{1}{similar\_sequences[s]}$
  **end for**
  **return** $sequence\_weight$
**end procedure**

##### S2.1.2 Sequence Weights

As described in the paper, NEFF can be considered as the normalized sum of weights for all sequences in an MSA. `NEFFy` offers the option to return the weight values of the sequences in the MSA instead of the final NEFF. This can be achieved by setting `only_weights` to `true`.

##### S2.1.3 Query Gaps

In some MSA files, such as the `sto` format, gaps may appear in the query sequence, indicating "gaps aligned to insertions". When calculating NEFF, users may prefer to exclude these gap positions from all sequences in the MSA. While most tools strictly adhere to the original MSA file, including any gaps in the query sequence, `NEFFy` provides flexibility in handling these gaps. It allows users to either filter them out, along with the corresponding positions in the aligned sequences (the default option), or retain them by setting `omit_query_gaps` to `false`.

It is also worth noting that the `Conkit` tool can handle gaps in the query sequence for the `a3m` format by offering two distinct options: `a3m-inserts` and `a3m`.

##### S2.1.4 Gappy Position

Introduced by `Gremlin`, gappy positions in a sequence alignment are those where the gap frequency exceeds a specific cutoff, indicating a higher-than-desired occurrence of gaps. Pseudo code is provided in algorithm 2. By filtering out these gappy positions, the NEFF calculation can focus on more informative and conserved regions of the sequences. The default approach in `NEFFy` includes all positions, treating none as gappy regardless of the number of gaps (`gap_cutoff = 1`). However, users can adjust the `gap_cutoff` parameter to any value greater than 0, up to 1, to manage gappy positions based on the specified cutoff.

##### S2.1.5 Alphabet

Each biological sequence is encoded using a valid set of characters to represent its composition. In the case of proteins, this set consists of 20 canonical amino acids, with each amino acid being

**Algorithm 2** Gappy Positions

```
procedure HandleGappyPositions(sequences, gap_cutoff)
    depth ← number of sequences
    gappy_positions ← []
    for each position p in alignment do
        gap_count[p] ← gaps in position p across sequences
        if gap_count[p] ≥ gap_cutoff × depth then
            append position p to gappy_positions
        end if
    end for
    for each sequence s in sequences do
        remove elements at positions listed in gappy_positions from sequence s
    end for
end procedure
```

represented by a specific letter. The list includes: ‘A’, ‘C’, ‘D’, ‘E’, ‘F’, ‘G’, ‘H’, ‘I’, ‘L’, ‘M’, ‘N’, ‘P’, ‘Q’, ‘R’, ‘S’, ‘T’, ‘V’, ‘W’, ‘Y’. Additionally, six non-standard amino acids, as detailed in section S2.1.6, also are included within the protein alphabet.

For DNA sequences, the alphabet comprises ‘A’, ‘T’, ‘C’, ‘G’ along with non-standard nucleic acid, ‘N’. Similarly, in the case of RNA sequences, the alphabet consists of ‘A’, ‘U’, ‘C’, ‘G’ along with non-standard nucleic acid ‘N’.

The default option in `NEFFy` is ‘protein’ alphabet.

##### S2.1.6 Non-Standard Residues

Non-standard residues refer to those that fall outside the typical set of residues, explicitly ‘N’ for DNAs and RNAs and ‘X’, ‘B’, ‘J’, ‘O’, ‘U’, ‘Z’ for proteins.

When calculating NEFF and determining sequence weights, various strategies can be applied for handling non-standard residues by configuring the `non_standard_option` parameter:

- **`AsStandard`**: Treat them as standard amino acids, following the approach of `DeepMSA`’s symmetric version.
- **`ConsiderGapInCutoff`**: To consider them as gaps only during sequence cutoff determination (the number of matches per sequence), aligning with `DeepMSA`’s asymmetric method.

- **ConsiderGap**: Treat them as gaps both in sequence cutoff calculation and when identifying matching positions between sequences, similar to the methods used by rMSA and Gremlin.

NEFFy is the only tool that supports all these options, with the default being AsStandard.

#### S2.2 Introduction of New Features

In addition to incorporating features from existing tools, NEFFy brings new capabilities for NEFF calculation, outlined in the subsections below.

##### S2.2.1 NEFF Calculation for Multiple MSAs

In all versions of NEFF calculation except for the NEFF of multimer MSA, NEFFy can accept multiple MSA input files as long as all sequences have the same length. It merges the sequences from these files in the specified order, removing any duplicates, and then computes the NEFF for the combined MSA.

##### S2.2.2 Per-Residue NEFF (Column-Wise NEFF)

This new feature computes NEFF for each position in the alignment. Per-residue NEFF values for each position in the MSA are calculated by summing the weights of the sequences that have a residue (i.e., non-gap characters) at that specific position. It is used by tools like AlphaFold [5] for more precise per-residue sequence diversity assessment.

##### S2.2.3 NEFF Calculation of Multimeric MSAs

The tool can calculate NEFF for MSAs of multimers, such as MSAs corresponding to multiple protein chains. Tools like AlphaFold-Multimer [3] generate these multimeric MSAs, where, for homomers (assemblies comprising multiple copies of the same chain), the MSA consists of multiple copies of the same sequence alignment. For heteromers (assemblies comprising two or more different chains), the MSA includes paired sequence alignments followed by individual MSAs for each chain in a block-diagonal arrangement. It's possible that some chains may not have an individual MSA, but a paired MSA is always present.

The tool can identify the multiple sequence alignment (MSA) format for both heteromers and homomers based on the provided stoichiometry of the complex. As described by the Protein Data Bank [11],"Stoichiometry indicates the number of chains participating in the assembly and whether the assembly is a homomer or a heteromer."

`NEFFy` computes NEFF values for such MSAs by using the stoichiometry and the lengths of the chains within the complex for heteromers. It identifies sequences that are either paired or unpaired across the chains, with the unpaired MSA sequences arranged in a block-diagonal pattern after the paired sequences. The tool then calculates NEFF for each of these components. For homomers, NEFF is calculated based on the individual MSA.

### S3 Exploring NEFF Calculation Tools

In this section, we will conduct a thorough exploration of the tools featured in Table 1 of the main paper which have integrated NEFF calculations into their core functions. We will delve into the primary purposes each of these tools serve and provide an in-depth exploration of the various features they offer for executing NEFF computations.

#### S3.1 `RaptorX`

`RaptorX`, a well-established protein structure prediction method developed since 2012, specializes in predicting protein tertiary and contact structures, particularly for sequences lacking close homologs in the Protein Data Bank [6]. `RaptorX` includes an integrated Python helper function for NEFF calculation, called **`Meff`**, which is specifically designed for computing asymmetric NEFF. This tool is primarily designed for handling aligned `a2m` and `a3m` formats, without the specification of biological sequence alphabets. The code is available on GitHub at `https://`

github.com/j3xugit/RaptorX-3DModeling/tree/master/BuildFeatures/Helpers.

#### S3.2 Conkit

Conkit is a Python library designed to facilitate the management and manipulation of residue-residue contact prediction data [12]. Among its key functionalities, it provides parsers for different MSA formats, allowing conversion between them, as well as the the capability to calculate NEFF and sequence weights, with a performance boost from Cython for time efficiency. The source code is available on GitHub at https://github.com/rigdenlab/conkit/blob/master/conkit/core/sequencefile.py, and the documentation can be found here.

Conkit does not specify the biological sequence type for NEFF computation. Additionally, when calculating symmetric NEFF, it gives an integer-valued NEFF value without normalization. Furthermore, Conkit can handle gaps in query sequences, but this option is available exclusively in the a3m format, distinguishing it from the a3m-inserts format, which includes gap positions of the query sequence in MSAs.

#### S3.3 DeepMSA

DeepMSA is an open-source tool designed for constriction of deep and sensitive MSAs [15]. It achieves this by leveraging homologous sequences and alignments derived from a diverse range of whole-genome and metagenome databases, utilizing complementary hidden Markov model algorithms. DeepMSA includes a built-in feature for NEFF computation in C++, which is exclusively available for the aln format. It offers support for both symmetric and asymmetric NEFF calculation for protein alphabets. The source code can be accessed on GitHub at https://github.com/kad-ecoli/MSAParser/blob/master/calNf.cpp.

##### S3.4 `Gremlin`

`Gremlin` is a method for prediction of residue-residue contacts that uses the power of sequence co-evolution and structural context data using a pseudo-likelihood approach. This allows for more precise prediction in protein structures, even when working with a limited set of homologous sequences [7]. `Gremlin` has been implemented in both Python and C++, and within its code, it includes the capability to compute symmetric NEFF.

When computing NEFF, `Gremlin` treats all non-standard residues as equivalent to gaps. It can support various biological sequences, including `protein`s, `RNA`s, and `DNA`s. `Gremlin` offers the feature to exclude gappy positions from the MSA and supports `fasta` and `aln` MSA formats. The source code is accessible on GitHub at `https://github.com/sokrypton/GREMLIN_CPP/blob/master/gremlin_cpp.cpp`.

##### S3.5 `rMSA`

`rMSA` is a hierarchical pipeline designed for conducting sensitive searches and precise alignments of RNA homologs for a RNA sequence [16]. `rMSA` includes a native feature for computing NEFF values specifically customized for RNA sequences. Research in RNA 3D structure prediction and protein-nucleotide structure prediction has employed `rMSA` to generate MSAs for RNA sequences [2, 4, 10]. The source code can be accessed on GitHub at `https://github.com/pylelab/rMSA/blob/master/src/fastNf.cpp`.

#### S4 Analysis and Results

##### S4.1 Reliability of `NEFFy` in Comparison with Other Tools

To evaluate the reliability and consistency of `NEFFy` compared to other tools, we calculated the NEFF value for each MSA file across 93 CASP15 targets. MSA files for these targets were generated using AlphaFold v2.3's default pipeline, which leverages three databases—Uniref90, Mg-

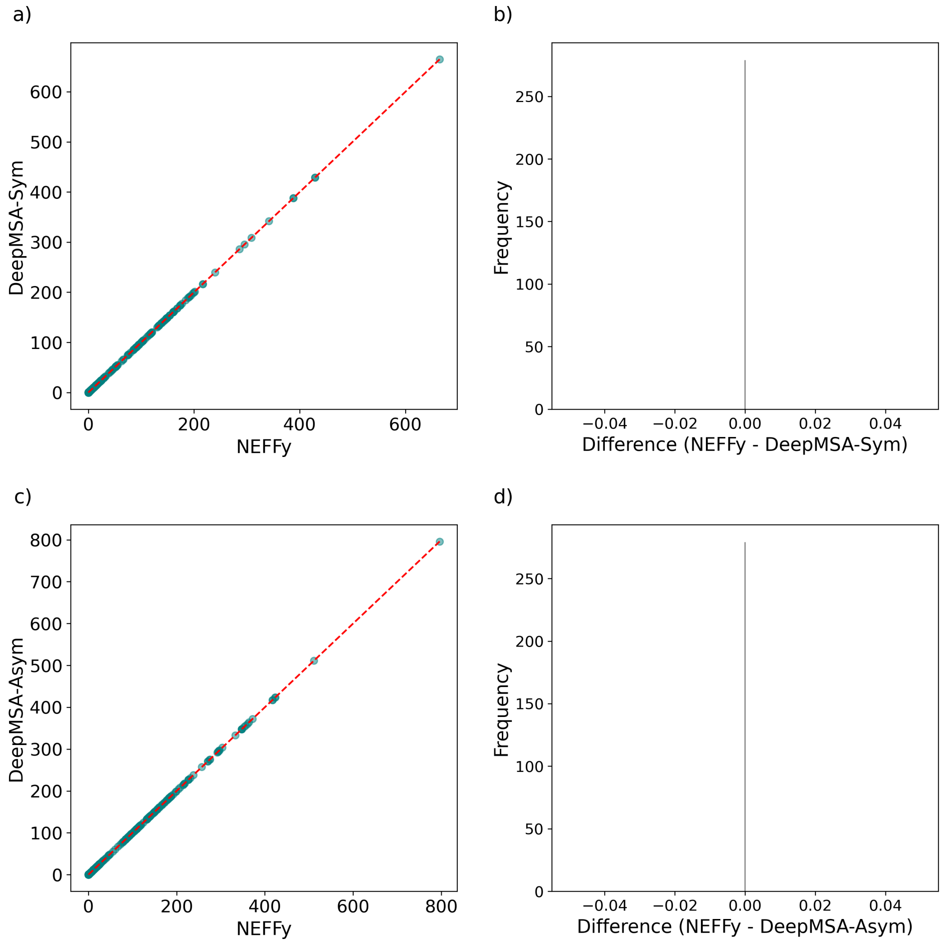

Figure S1: Comparison of NEFF values between `DeepMSA` and `NEFFy` (using parameters corresponding to `DeepMSA`): (**a**, **c**) Scatter plots showing NEFF values with a diagonal reference line for symmetric (**a**) and asymmetric (**c**) versions. (**b**, **d**) Histograms showing the distribution of differences for symmetric (**b**) and asymmetric (**d**) versions.

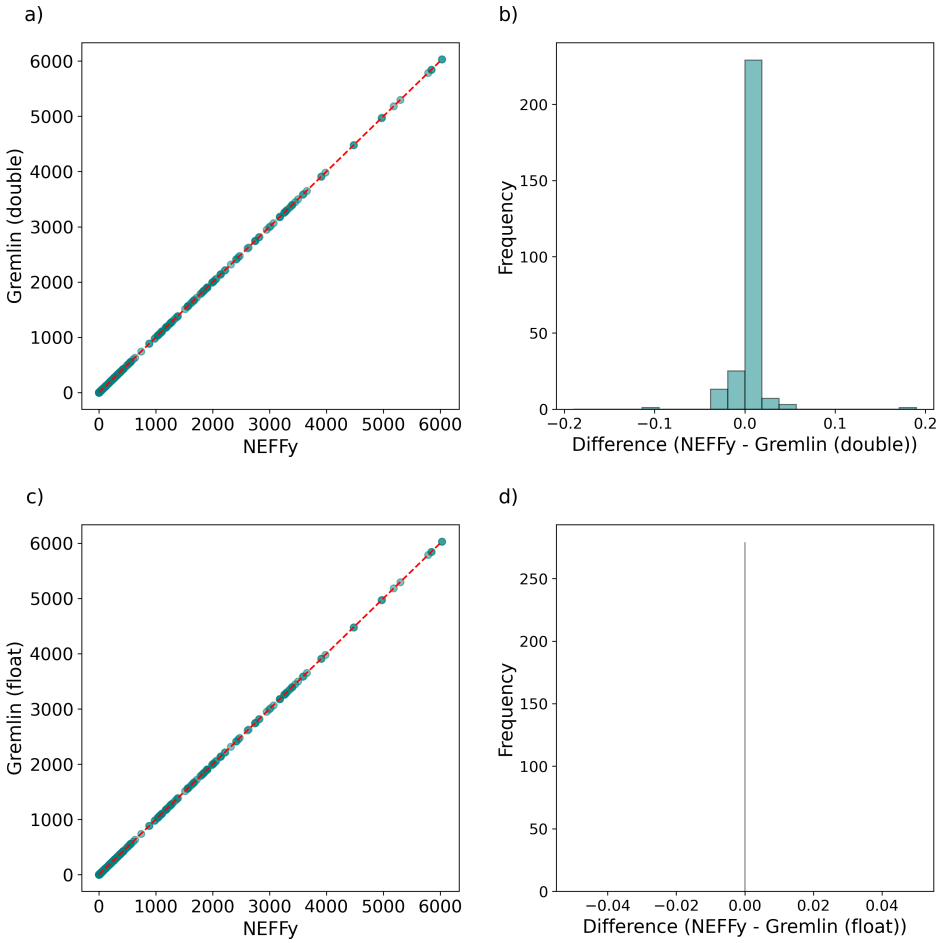

Figure S2: Comparison of NEFF values between `Gremlin`, using `double` (default version) and `float` data types, and `NEFFy` (using parameters corresponding to `Gremlin`) for the CASP15 dataset: (**a**, **c**) Scatter plots showing NEFF values with a diagonal reference line for `double` (**a**) and `float` (**c**) configurations. (**b**, **d**) Histograms showing the distribution of differences for `double` (**b**) and `float` (**d**) configurations.

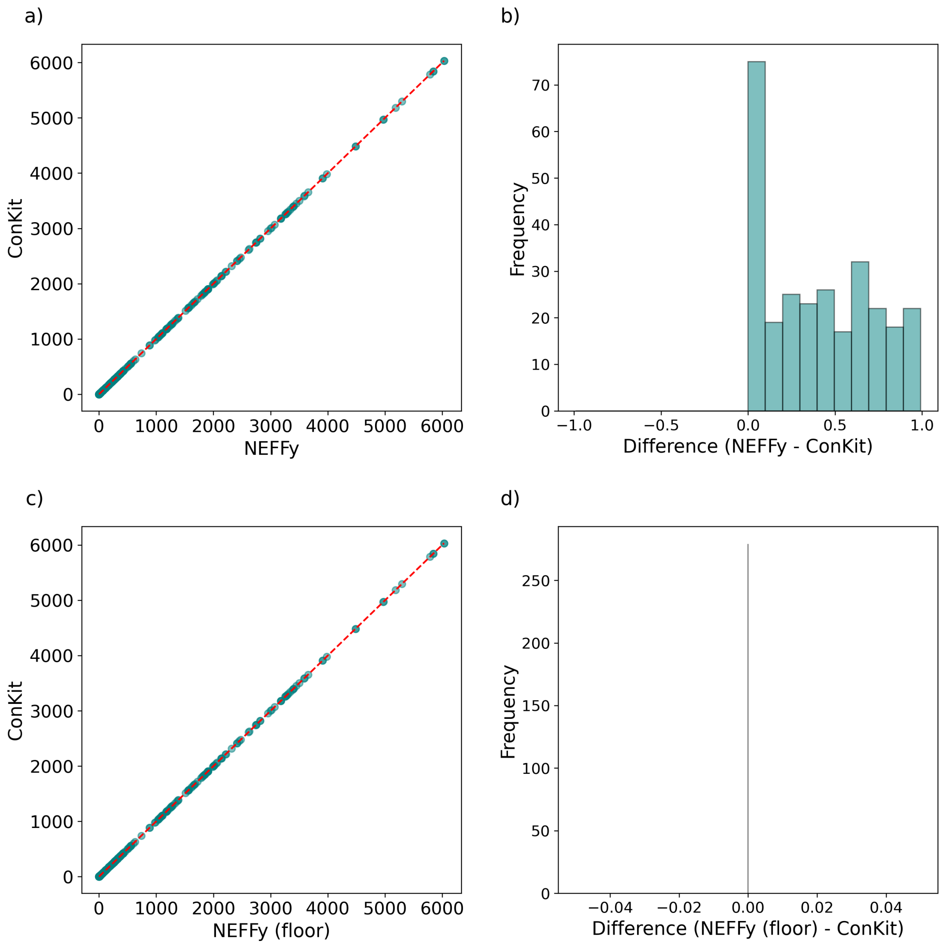

Figure S3: Comparison of NEFF values between `ConKit` and `NEFFy`, evaluated with both default and 'rounded NEFF values' configurations (using parameters aligned with `ConKit`) for the CASP15 dataset: (**a**, **c**) Scatter plots showing NEFF values with a diagonal reference line for default (**a**) and 'rounded NEFF values' (**c**) configurations. (**b**, **d**) Histograms showing the distribution of differences for default (**b**) and 'rounded NEFF values' (**d**) configurations.

| Tool | Mode | Commands |
|---|---|---|
| DeepMSA | Symmetric | DeepMSA: **./calNf [file] 0.8 0**<br>NEFFy: **./neff --file=[file]** |
| | Asymmetric | DeepMSA: **./calNf [file] 0.8 10**<br>NEFFy: **./neff --file=[file] --is_symmetric=false --non_standard_option=1** |
| Gremlin | | Gremlin: **./gremlin_cpp -i [file] -eff_cutoff 0.8 -gap_cutoff 1 -only_neff**<br>NEFFy: **./neff --file=[file] --norm=2 --non_standard_option=2** |
| Conkit | | Conkit:<br>import conkit.io<br>msa = conkit.io.read(file, format)<br>neff = msa.meff<br>depth = msa.nseq<br><br>NEFFy: **./neff --file=[file] --norm=2** |
| RaptorX | | RaptorX: run *RaptorX* python source with threshold = 0.8<br>NEFFy: **./neff --file=[file] --is_symmetric=false --norm=2** |
| rMSA | | rMSA: **./fastNf [file] 0.8 0**<br>NEFFy: **./neff --file=[file] --non_standard_option=2** |

Table S2: Commands for calculating NEFF values with each tool and their equivalents when using NEFFy.

nify, and BFD—resulting in three separate MSA files per target, totaling 279 files in both STO and A3M formats. Since some tools do not support NEFF calculation in these formats, we used NEFFy's built-in converter to transform the files into compatible formats for each tool (available at https://zenodo.org/records/14210950). Table S2 provides a summary of the commands required to calculate NEFF values with each tool, including both the commands using the tool and their equivalents when using NEFFy.

Our findings demonstrated complete consistency with both the symmetric and asymmetric options of DeepMSA (Figure S1). For Gremlin, only minor differences were observed due to its use of the double data type, while NEFFy and DeepMSA use float for decimal representation. These differences in data types impact precision and storage size, which can cause slight variations in results during mathematical operations. However, when the data type in Gremlin was changed to float, the results matched entirely (Figure S2). In the case of Conkit, the tool rounds NEFF values to integers, which explains the subtle differences observed in NEFF results. When the NEFF values from NEFFy were rounded, the results aligned with those of Conkit (Figure S3). For RaptorX, we observed slight variations, primarily due to its consistent exclusion of lowercase residues from sequences and its unique method for determining cutoffs by taking the minimum length of each sequence pair during similarity calculations. Additionally, the imple-

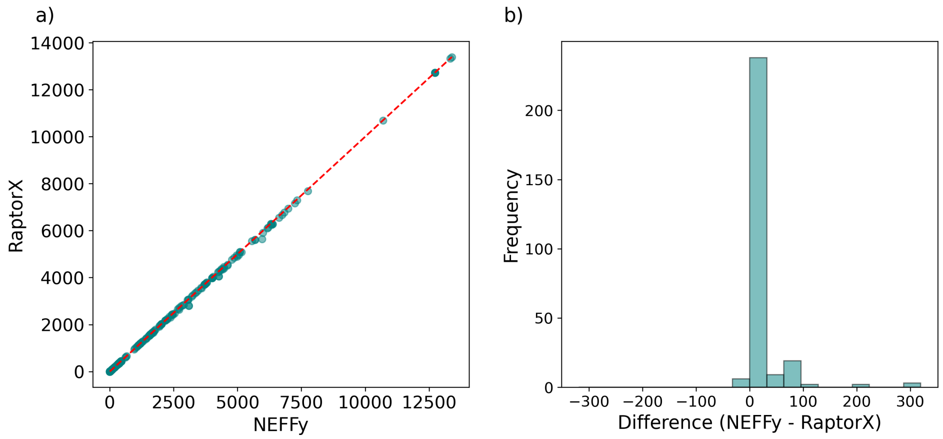

Figure S4: Comparison of NEFF values between `RaptorX` and `NEFFy` (using parameters corresponding to `RaptorX`). (**a**) Scatter plot showing NEFF values with a diagonal reference line. (**b**) Histogram showing the distribution of differences.

mentation in Python may introduce minor discrepancies in handling decimal numbers (Figure S4).

Additionally, for `rMSA`, we used the benchmark dataset described in [16], which includes 361 non-redundant RNA chains collected from the PDB database. We generated MSAs for these chains using `rMSA` tool (available at `https://zenodo.org/records/14210950`). We then conducted a comparative analysis by comparing our NEFF values with those generated by the built-in `rMSA` NEFF computation tool for these files. The `NEFFy` results showed minor differences compared to the built-in tool, as `NEFFy` uses the float data type while `rMSA` uses double. As a test, we changed `rMSA`'s data type to float and obtained identical results (Figure S5).

#### S4.2 Computational Efficiency of `NEFFy`

We used the same set of 279 MSA files generated by AlphaFold for CASP15 targets. Each tool was run on these files five times to record execution times, and the average execution time for each file and tool was calculated. Figure S6 illustrates the distribution of execution times across the various tools.

`RaptorX`, implemented entirely in Python, utilizes nested loops for its similarity calculations. This approach can be inefficient, particularly with large datasets, resulting in significant perfor-

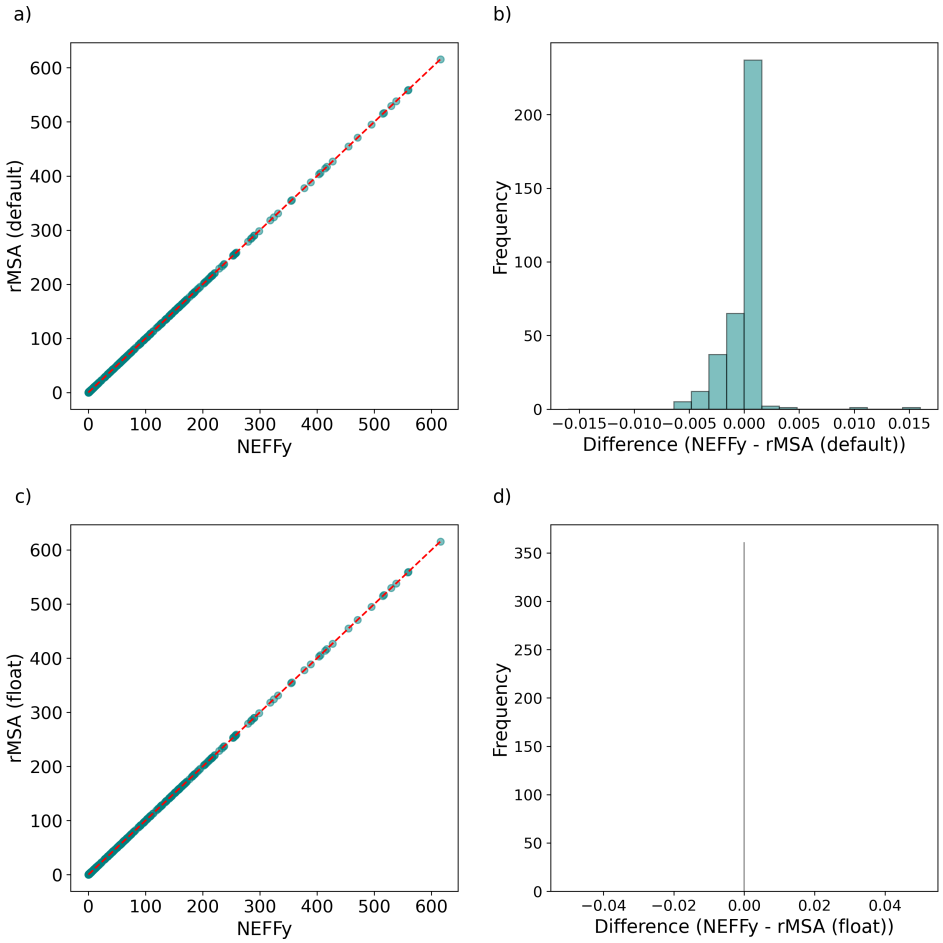

Figure S5: Comparison of NEFF values between `rMSA`, using `double` (default version) and `float` data types, and `NEFFy` (using parameters corresponding to `rMSA`) for the `rMSA` dataset: (**a**, **c**) Scatter plots showing NEFF values with a diagonal reference line for `double` (**a**) and `float` (**c**) configurations. (**b**, **d**) Histograms showing the distribution of differences for `double` (**b**) and `float` (**d**) configurations.

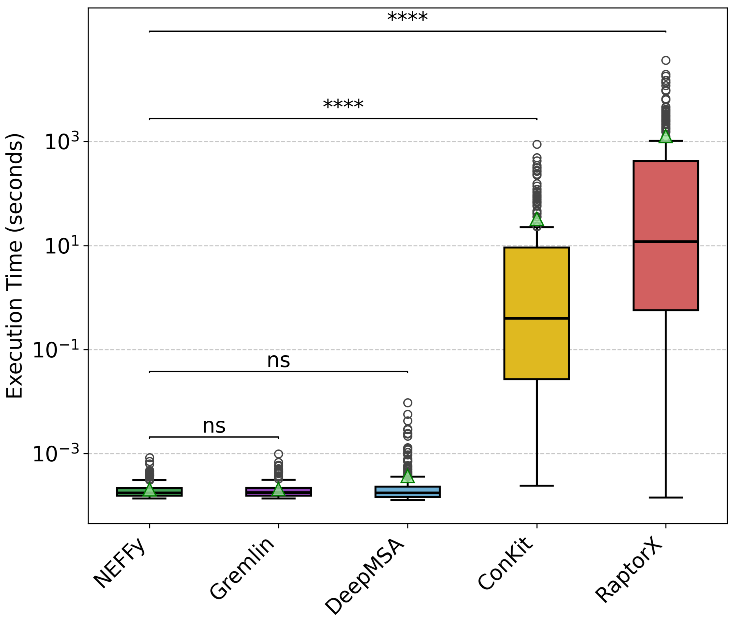

Figure S6: Comparison of execution times (in seconds) for the tools `NEFFy`, `DeepMSA`, `Gremlin`, `Conkit`, and `RaptorX` on a log scale, based on the average from five runs. The mean execution times for each tool are indicated by green triangles. Statistical significance was calculated using one-sided Wilcoxon test, with non-significant differences denoted as “ns” and significant differences represented by asterisks (****).

mance bottlenecks and slower processing speeds. In contrast, `Conkit`, also written in Python, leverages Cython code segments to enhance its performance through C-like syntax and optimizations. Conversely, `DeepMSA`, `Gremlin`, `rMSA`, and `NEFFy` are considerably faster due to their implementation in C++, which allows for compiler optimizations that improve processing speed and overall performance.

Statistical analysis using the one-sided Wilcoxon signed-rank test [13] further supports these performance differences. `NEFFy` demonstrates comparable performance to `DeepMSA` ($p \approx 0.027$) and `Gremlin` ($p \approx 0.081$), showing no significant difference. However, `NEFFy` significantly outperforms `Conkit` ($p \approx 8.29 \times 10^{-48}$) and `RaptorX` ($p \approx 8.94 \times 10^{-48}$).

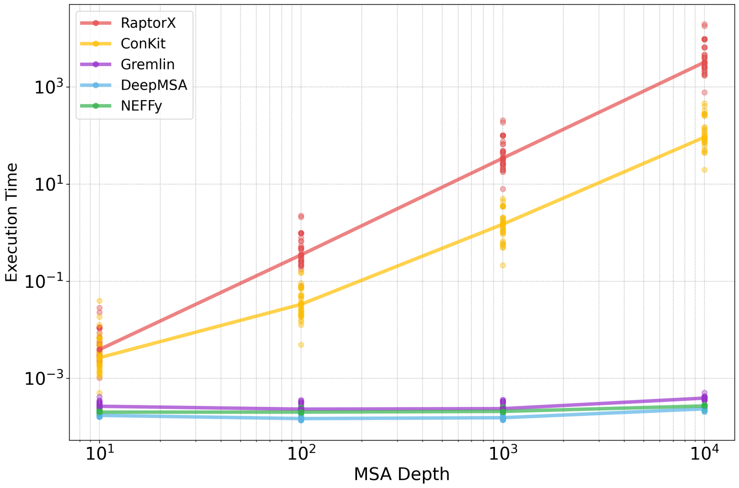

Figure S7: Scalability analysis of five tools—`NEFFy`, `DeepMSA`, `Gremlin`, `Conkit`, and `RaptorX`—based on execution time versus MSA depth for 36 MSA files, each corresponding to a target from CASP15. The plot demonstrates how each tool's performance changes with increasing MSA depth. Execution times were measured across depths from 10 to 10,000, with individual points representing the average execution time (from five runs) per file at each depth. Solid lines show the median execution time across files for each tool.

#### S4.3 Scalability Assessment

We used 36 targets from CASP15, each with a UniRef90 MSA of 10,000 sequences, to evaluate NEFF computation for all tools across four depths: 10, 100, 1000, and 10,000. `NEFFy` supports custom depths directly, while for other tools, we used the complete MSA for a depth of 10,000 and extracted the first 10, 100, and 1,000 sequences from the MSA file to calculate NEFF for depths of 10, 100, and 1,000, respectively. We measured the execution time for each tool and depth across five separate runs and calculated the average time for each target from these five executions to evaluate scalability as depth increased.

Fig. S7 demonstrates the scalability of five different methods—`NEFFy`, `DeepMSA`, `Gremlin`, `Conkit`, and `RaptorX`—as a function of increasing MSA depth on a logarithmic scale for both depth and execution time. `NEFFy`, `DeepMSA`, and `Gremlin` maintain consistent performance across all tested depths, indicating that these tools are well-optimized and highly scalable for both shallow and deep MSAs, making them particularly suited for large-scale analyses where computational efficiency is critical. In contrast, `Conkit` and `RaptorX` exhibit a noticeable increase in

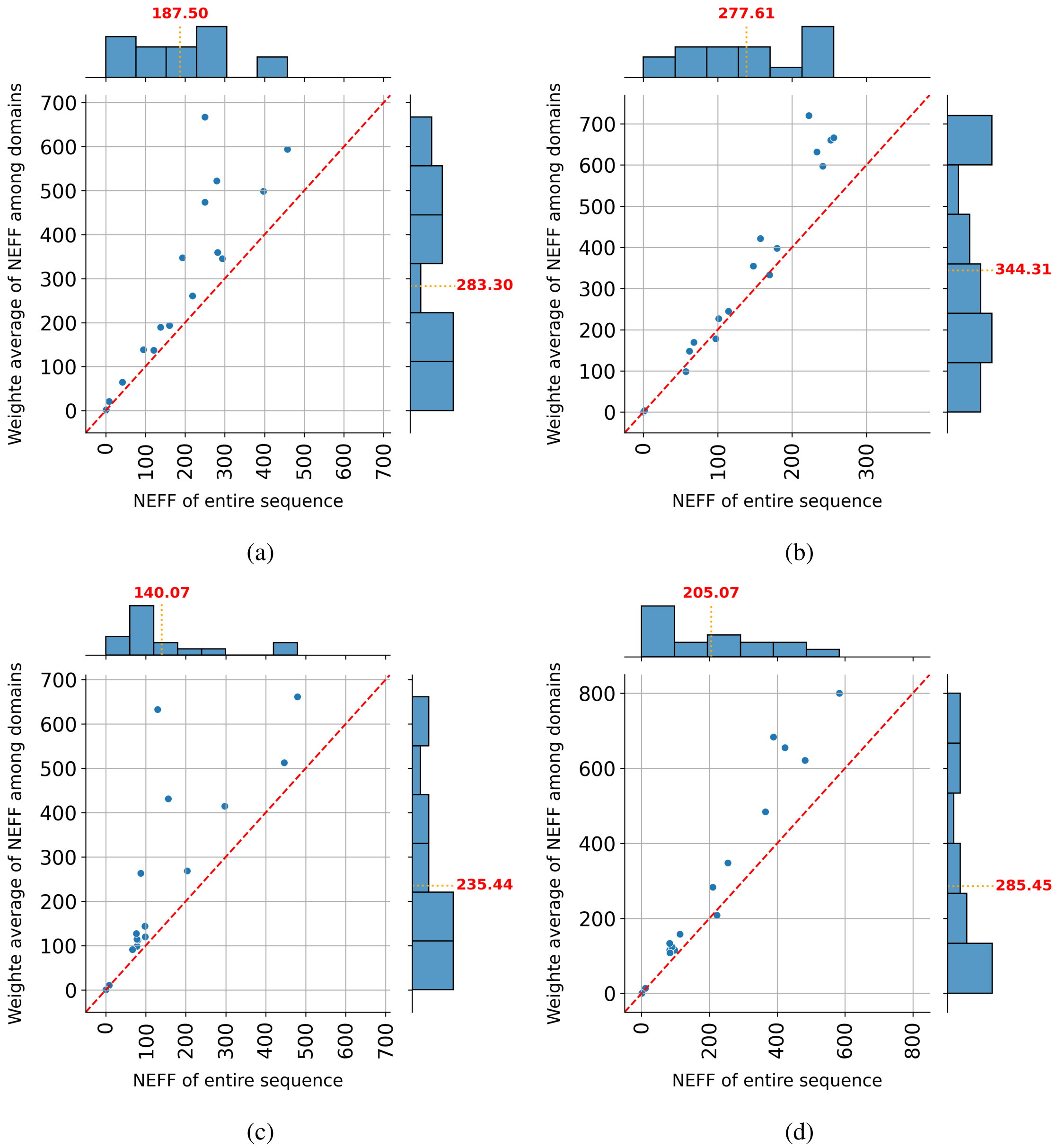

Figure S8: Domain-level NEFF vs. whole target for 17 multi-domain targets in CASP15: **a, b)** AlphaFold: The default pipeline for MSA generation was used, combining three distinct MSAs per target to produce the final MSA. **(a)** Symmetric NEFF **(b)** Asymmetric NEFF **c, d)** RoseTTAFold: The default MSA generation pipeline was employed. **(c)** Symmetric NEFF **(d)** Asymmetric NEFF.
All analyses used normalized NEFF values (sqrt(L)).

execution time with growing depth, following an exponential pattern, which suggests these tools may struggle to scale effectively with deeper MSAs, rendering them less practical for extensive MSA applications compared to the other tools.

#### S4.4 Case Study on Multi-domain Proteins

Following our analysis of the 19 multidomain sets from the CASP15 dataset, we utilized the target MSAs generated by AlphaFold and RoseTTAFold [1], as provided in the paper by Moussad et al. at `https://zenodo.org/records/7682977`. Since RoseTTAFold generates a single MSA file, we were able to use the provided MSAs directly without any additional steps. For AlphaFold, we used the default AlphaFold 2.3 pipeline, which merges the three MSAs before feeding them into the deep learning model. The combined MSA was subsequently used for our analysis. To obtain domain-level MSAs, we extracted the sequence alignments specific to each domain from the full-length MSA of the target, as detailed in [14]. We then used `NEFFy` to calculate normalized NEFF values (`sqrt of L`) for both the full protein and domain-level MSAs. To getting the NEFF values for individual domains, we utilized the `start_pos` and `end_pos` features of `NEFFy`, setting the domain's start and end positions accordingly. For two of the targets, "T1157s1" and "T1158," NEFF values could not be obtained because one of their domains was fragmented across multiple locations. Therefore, we proceeded with our analysis on 17 such targets.

Fig. S8 illustrates the Grishin plots [8], which present the weighted average NEFF values for the extracted MSAs of individual domains alongside the NEFF values for the MSAs of the entire protein chains. These plots illustrate the correlation between the weighted sum of the NEFF values for the domains (on the y-axis) and the NEFF values for the entire protein chain (on the x-axis), indicating that the NEFF values of individual domain MSAs are higher than the NEFF values of entire chain MSAs. The observed pattern aligns with the trends noted in prediction accuracy [9].
